# Clemastine fumarate enhances myelination and promotes functional recovery in a syndromic ASD mouse model of Pitt-Hopkins Syndrome

**DOI:** 10.1101/2022.05.03.490512

**Authors:** Joseph F. Bohlen, Colin M. Cleary, Debamitra Das, Srinidhi Rao Sripathy, Norah L. Sadowski, Gina Shim, Rakaia F. Kenney, Ingrid P. Buchler, Daniel K. Mulkey, Brady J. Maher

## Abstract

Pitt-Hopkins syndrome (PTHS) is an autism spectrum disorder (ASD) caused by autosomal dominant mutations in the Transcription Factor 4 gene (*TCF4*). One pathobiological process caused by *Tcf4* mutation is a cell autonomous reduction in oligodendrocytes (OLs) and myelination. In this study, we show that clemastine is effective at restoring myelination defects in a PTHS mouse model. In vitro, clemastine treatment reduced excess oligodendrocyte precursor cells (OPCs) and normalized OL density. In vivo, two-week intraperitoneal administration of clemastine also normalized OPC and OL density in the cortex of *Tcf4* mutant mice and appeared to increase the number of axons undergoing myelination, as EM imaging of the corpus callosum showed a significant increase in uncompacted myelin. Importantly, this treatment paradigm resulted in functional rescue by improving electrophysiology and behavior. Together, these results provide preclinical evidence that remyelination therapies may be beneficial in PTHS and potentially other neurodevelopmental disorders characterized by demyelination.

## Introduction

Autism spectrum disorder (ASD) is genetically heterogeneous with convergent symptomatology, suggesting the potential for common dysregulated pathways. Currently, there are no therapeutic interventions that address the core symptoms of ASD. One form of syndromic ASD caused by autosomal dominant mutations in the transcription factor 4 (*TCF4;* not *TCF7L2 /*T-Cell Factor 4) gene results in Pitt-Hopkins syndrome (PTHS), a rare neurodevelopmental disorder characterized by intellectual disability, failure to acquire language, deficits in motor learning, hyperventilation, gastrointestinal abnormalities, and autistic behavior (1). Mouse models of PTHS consistently show behavioral deficits that approximate behavioral abnormalities observed in PTHS patients (2–5). However, the pathophysiological mechanisms underlying these behavioral deficits are not completely understood.

Transcriptional profiling across 5 independent PTHS mouse models identified enrichment in differentially expressed genes (DEGs) related to myelination. This transcriptional profile was biologically validated, whereby in vivo, in vitro, and ex vivo experiments demonstrated mutations in *Tcf4* resulted in a reduction in oligodendrocytes (OLs) in conjunction with demyelination related functional deficits (6). *Tcf4* is highly expressed in the entire OL lineage, and reductions in myelination due to *Tcf4* mutations are cell autonomous (6–10). Beyond PTHS, defects in the OL lineage are reported for a variety of ASD models and suggests this cell population may be a suitable target for therapeutic interventions. One well known remyelination compound is clemastine fumarate, which is a well-studied, FDA-approved, first-generation antihistamine predominantly used in the treatment of allergic conditions (11). It is a competitive antagonist at peripheral H1 receptors, blocking the actions of endogenous histamines, but its remyelination properties are thought to be through its muscarinic receptor 1 (M1R) activity (12–14). Several studies have consistently shown clemastine promotes differentiation of OPCs into mature/myelinating oligodendrocytes (13,15–17). Remyelination is an intriguing therapeutic approach for neurodevelopmental disorders (NDDs) as the presence of mitotically active OPCs and the process of myelination occurs over the entire lifespan (18). In this study, we demonstrate that pharmacological enhancement of myelination with clemastine is a beneficial rescue approach in a PTHS mouse model. We show that clemastine administration, both in vitro and in vivo, normalizes OPC and OL density, improves myelination, and normalizes electrophysiological and behavioral deficits.

## Methods

### Animals and tissue collection

The *Tcf4*^*+/tr*^ mouse model of PTHS is heterozygous for an allele encoding deletion of the DNA-binding domain of TCF4 (B6;129-TCF4tm1Zhu/J, stock number 013598, Jackson Laboratory). This mouse colony was backcrossed for at least six generations, maintained by The Lieber Institute for Brain Developments Animal Facility on a 12-h light/dark cycle and fed ad libitum. *Tcf4*^*+/tr*^ mouse samples were matched with samples from *Tcf4*^*+/+*^ littermates, and sex was randomly selected in each genotype and age group. All procedures were performed in accordance with the National Institutes of Health Guide for the Care and Use of Laboratory Animals and approved by the Johns Hopkins University School of Medicine’s Institutional Animal Care and Use Committee.

### Clemastine Dosing Regime

A dosing solution at concentration 1 mg/mL was generated by weighing out the appropriate quantity of clemastine fumarate in a septa seal vial. 5% DMA (dimethylacetamide, Sigma 271012) was added up to the total volume needed. The clemastine fumarate was completely dissolved in the DMA prior to the addition of saline to the appropriate volume. Vehicle was consisting of 5% DMA in phosphate buffered saline. The mixture was shaken vigorously (vortex) and was ready for administration for intraperitoneal (IP) dosing at 10 mL/kg. The final pH was approximately 5. Animals were dosed every 24 hours for 14 consecutive days at a dose of 10mg/kg for clemastine or an equal volume of vehicle depending on the condition. Initial dosing was performed at P28 up until P42 when animals were then used for subsequent experiments.

### Primary OPC and OL cultures

Primary OPC and OLs were obtained following a previous protocol (19). In brief, P2-P3 pups were dissociated and plated at a density of 10.0 × 10^3^ cells/cm^2^ in a 96 well ibidi plate coated in 0.1% polyethleneimine (PEI) and 0.5 μg/mL laminin. Cells were plated and maintained in OPC proliferation media consisting of 1x StemPro Neural supplement (Thermo Fisher Scientific, A1050801), 1x Anti-Anti (Thermo Fisher Scientific, 15240-096), 10 ng/mL of Human FGF-basic (Peprotech, 100-18B) and 30ng/mL rhPDGF-AA (R&D systems, 221-AA), with a half media exchange on DIV4. On DIV7 OL differentiation media was added (base media with removal of bFGF and rhPDGF-AA) and cells were differentiated into oligodendrocytes with the addition of either DMSO (Sigma D8418 at 0.1%) as a vehicle or Clemastine (Sigma SML0445 10μM) with media change every day until DIV14.

### Immunohistochemistry and Immunocytochemistry

Cells were rinsed 1x with PBS and then fixed with 4% paraformaldehyde (PFA) for 5 minutes, following fixation cells were rinsed 3x with PBS. Similarly, mice were perfused with 20 - 30mLs of 1x PBS, followed by 20 - 30mLs of 4% PFA. Tissue was extracted and post fixed in 4% PFA on a rocker at 4°C overnight. For immunostaining cells were rinsed 3x with 0.04% Tween 20 while tissue (P42 mouse) was rinsed 3x with 0.4% Triton. Cells were blocked in respective serum (10%) for 2 hours at room temperature on an orbital shaker. Following block, primary antibody was added in 2% serum in 0.04% Tween 20 for cells and 0.4% Triton-X 100 for tissue, and incubated overnight at 4°C. Following overnight incubation cells were rinsed 3x with 0.04% Tween 20 or 0.4% Triton respectively before adding the secondary antibody to incubate at room temperature for 2 hours. After incubation, cells were rinsed 3x in respective buffers and counterstained with DAPI (Invitrogen™, D1306). Visualization was carried out on a ZEISS LSM 700 Confocal. Imaging and quantification were performed blind to genotypes and conditions/treatments.

### Transmission Electron Microscopy (TEM)

After perfusion with a 0.1 M sodium cacodylate buffer, pH7.2, containing 2% paraformaldehyde (freshly prepared from EM grade aqueous solution), 2% glutaraldehyde, and 3mM MgCl2, p42 mouse brains were kept overnight in fixative. The next day brains were dissected in fixative and rinsed with sodium cacodylate buffer. Samples were then post-fixed in reduced 2% osmium tetroxide, 1.6% potassium ferrocyanide in buffer (2 hr) on ice in the dark. Following a dH2O rinse, samples were stained with 2% aqueous uranyl acetate (0.22 µm filtered, 1 hr, dark), dehydrated in a graded series of ethanol, propylene oxide and embedded in Eponate 12 (Ted Pella) resin. Samples were polymerized at 60°C overnight. Thin sections, 60 to 90 nm, were cut with a diamond knife on the Reichert-Jung Ultracut E ultramicrotome and picked up with copper slot (1 × 2 mm) grids. Grids were stained with 2% uranyl acetate and observed with a Phillips CM120 TEM at 80kV. Images were captured with an AMT XR80 CCD camera. Preparation of samples, TEM imaging, and quantification was performed blind to genotypes.

### Electrophysiology

Acute coronal brain slices containing the corpus callosum (CC) were obtained from P39–P42 mice as previously described (20). Artificial cerebrospinal fluid (ACSF) was oxygenated (95% O2 and 5% CO2) and contained (in mM): 125 NaCl, 25 NaHCO3, 1.25 NaH2PO4, 3 KCl, 25 dextrose, 1 MgCl2, and 2 CaCl2, pH 7.3. A bipolar stimulating electrode was placed 500µm away from the midline and CC was stimulated with a 100µs square pulse using 80%of the maximal stimulation intensity. The recording electrodes were fabricated from borosilicate glass (N51A, King Precision Glass, Inc.) to a resistance of 2–5 MΩ and placed at varying distances from the stimulating electrode in the contralateral CC. For cAP recording, pipettes were filled with ACSF. Voltage signals were recorded with an Axopatch 200B amplifier (Molecular Devices) and were filtered at 2 kHz using a built in Bessel filter and digitized at 10 kHz. Data was acquired using Axograph on a Dell PC. For electrophysiology experiments, data collection and analysis were performed blind to the conditions of the experiment.

### Novel Open field assay

Locomotor activity and anxiety was assessed using Noldus PhenoTyper cages as previously shown (21). Each cage is outfitted with two sets of cameras; one on the ceiling that faces the platform (35 cm × 35 cm), and another pointed at the side of the cage. Mice were acclimated to the experimentation room in their home cages for at least 1 h. During acclimation, the Noldus EthoVision software was set-up to track movement for 30 total minutes. After acclimation, mice were placed in the center of the open field, opaque Plexiglas was placed on all four sides of the cage to obscure any visual cues, and the trial was started in the EthoVision software. After 30 min, the trial ended, and mice were placed back into their home cages. Noldus EthoVision software was used to determine distance traveled, time spent in center, and frequency of going to center.

### Statistics

GraphPad Prism (GraphPad Software, San Diego, CA) was used to conduct statistical analyses for IHC, ICC, and behavioral experiments. Scipy stats package version 1.8.0 and Statsmodels package version 0.13.2 were used to conduct statistical analyses for EM and electrophysiology experiments. Data were analyzed using either a two-way analysis of variance (ANOVA), analysis of covariance (ANCOVA), or an unpaired t test. All ANOVA main effects were followed by Tukey post hoc tests.

## Results

### Clemastine induces maturation of OPCs in vitro

We first tested the effects of clemastine administration on maturation of OPCs in vitro. Previous studies indicate that clemastine is effective at promoting OPCs in primary cultures to differentiate into mature OLs (MBP+) (12,19). Following a previously established protocol (19), we dissociated and plated OPCs from both *Tcf4*^*+/+*^ and *Tcf4*^*+/tr*^ mice onto 96 well plates in OPC proliferation media. OPCs were differentiated on DIV7 with OL differentiation media containing either clemastine (1μM) or vehicle (DMSO) and fed every day with fresh media containing clemastine or vehicle before immunostaining on DIV14 (Figure 1A). We performed blinded quantification of OPCs and OLs using antibodies against PDGFRα and MBP, respectively, and normalized our counts with the pan-OL marker OLIG2 (Figure 1A). Consistent with previous observations (6), *Tcf4*^*+/tr*^ cultures treated with vehicle exhibited a significant decrease in mature OLs (MBP+/Olig2+) along with a significant increase in OPCs (PDGFRα+/Olig2+) when compared to littermate controls (Figure 1C, D). Clemastine treatment resulted in a significant reduction in OPCs and an increase in OLs when compared to vehicle treated *Tcf4*^*+/tr*^ cells (Figure 1C-D). These data indicate clemastine is effective at overcoming mutations in Tcf4 by promoting differentiation of OPCs into mature OLs and thereby normalizing the OL population to similar proportions of OPCs and OLs observed in *Tcf4*^*+/+*^ cultures.

**Figure 1.**
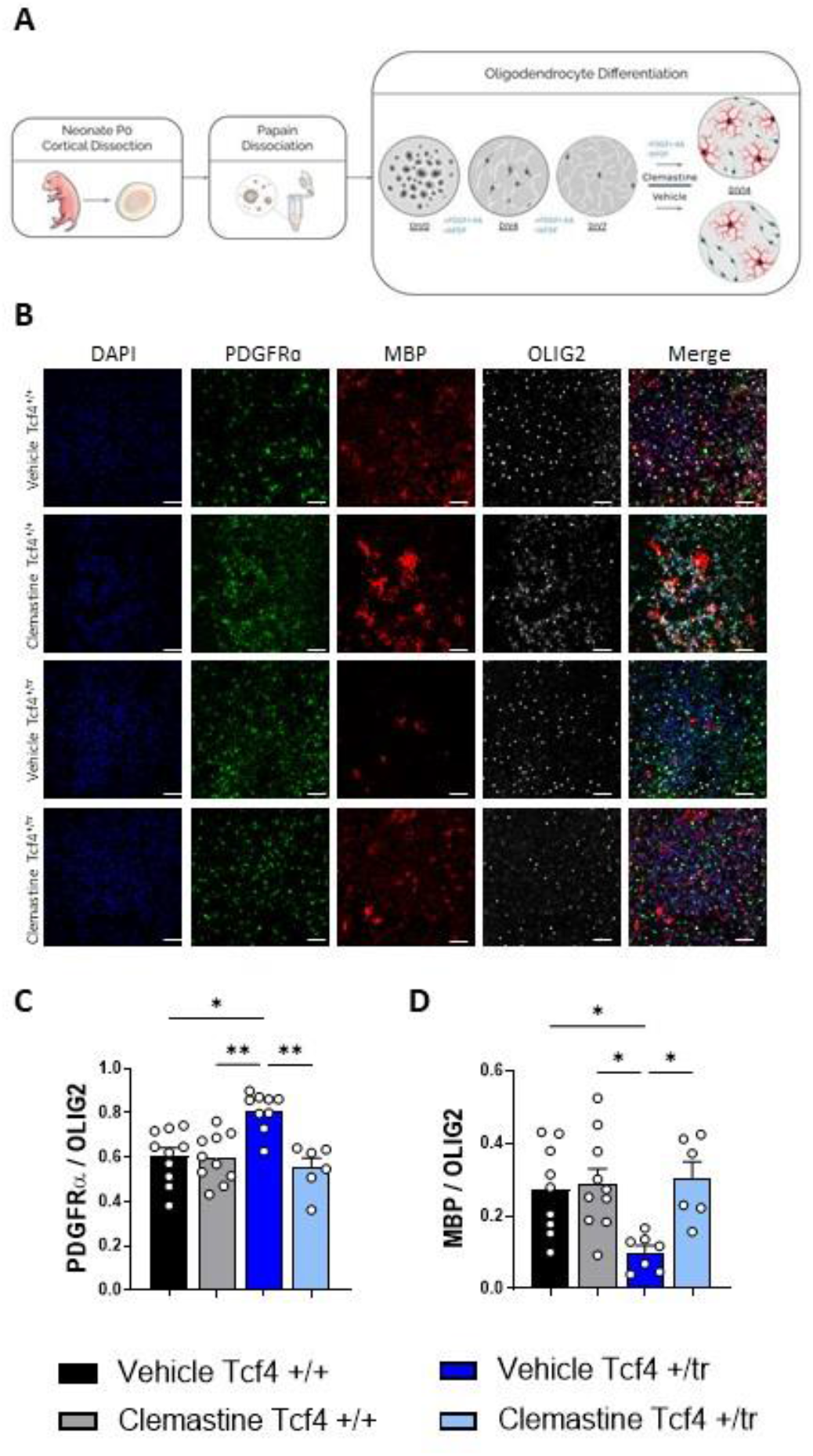
Clemastine increases maturation of oligodendrocytes in *Tcf4*^*+/tr*^ cultures. **(A)** Graphical representation of plating and dosing regimen. **(B)** Immunocytochemistry of OPC/OL DIV14 in culture following 7 days of either vehicle (DMSO) or clemastine (1μM), cultures stained for DAPI (blue), PDGFRα (green), MBP (red) and Olig2 (gray). **(C-D)** Summary plots show *Tcf4*^*+/+*^ cells with vehicle (black) or treated with clemastine fumarate (gray) as well as *Tcf4*^*+/tr*^ cells with vehicle (dark blue) or treated with clemastine fumarate (sky blue). Representative results for response to vehicle and clemastine administration comparing PDGFRα/Olig2 (immature population) (Two-way ANOVA n=10-13 biologically independent animals/genotype F_3,22_=14.39 p<0.0001 data are presented as mean values ± SEM) and MBP+/Olig2+ (Two-way ANOVA n=11-13 biologically independent animals/genotype F_3,19_=5.966 p=0.0048 data are presented as mean values ± SEM). These data show clemastine fumarate increased the percentage of mature OLs (MBP) while reducing the overall pool of OPCs (PDGFRα) in vitro. *p<0.05, **p<0.01, ***p<0.001, ****p<0.0001.

### Clemastine increases the number of CC-1 positive cells in vivo

Next, we assessed the effectiveness of clemastine in vivo by dosing *Tcf4*^*+/tr*^ mice and *Tcf4*^*+/+*^ littermates intraperitoneally with either clemastine (10 mg/kg) or vehicle for two weeks followed by immunohistochemical (IHC) quantification (Figure 2A). Consistent with prior results (6), vehicle-treated *Tcf4*^*+/tr*^ mice showed fewer mature OLs (CC1+/Olig2+) and more OPCs (PDGFRα+/Olig2+) when compared to vehicle-treated *Tcf4*^*+/+*^ littermates (Figure 2C, D). Similar to our in vitro data, clemastine-treated *Tcf4*^*+/tr*^ mice showed a significant increase in mature OLs that coincided with a significant decrease in OPCs when compared to their vehicle-treated *Tcf4*^*+/+*^ littermates (Figure 2C, D). Together, these data indicate clemastine is also effective in vivo at promoting OPCs to differentiate into OLs in a mutant Tcf4 background, thereby normalizing the OL population to *Tcf4*^*+/+*^ levels.

**Figure 2.**
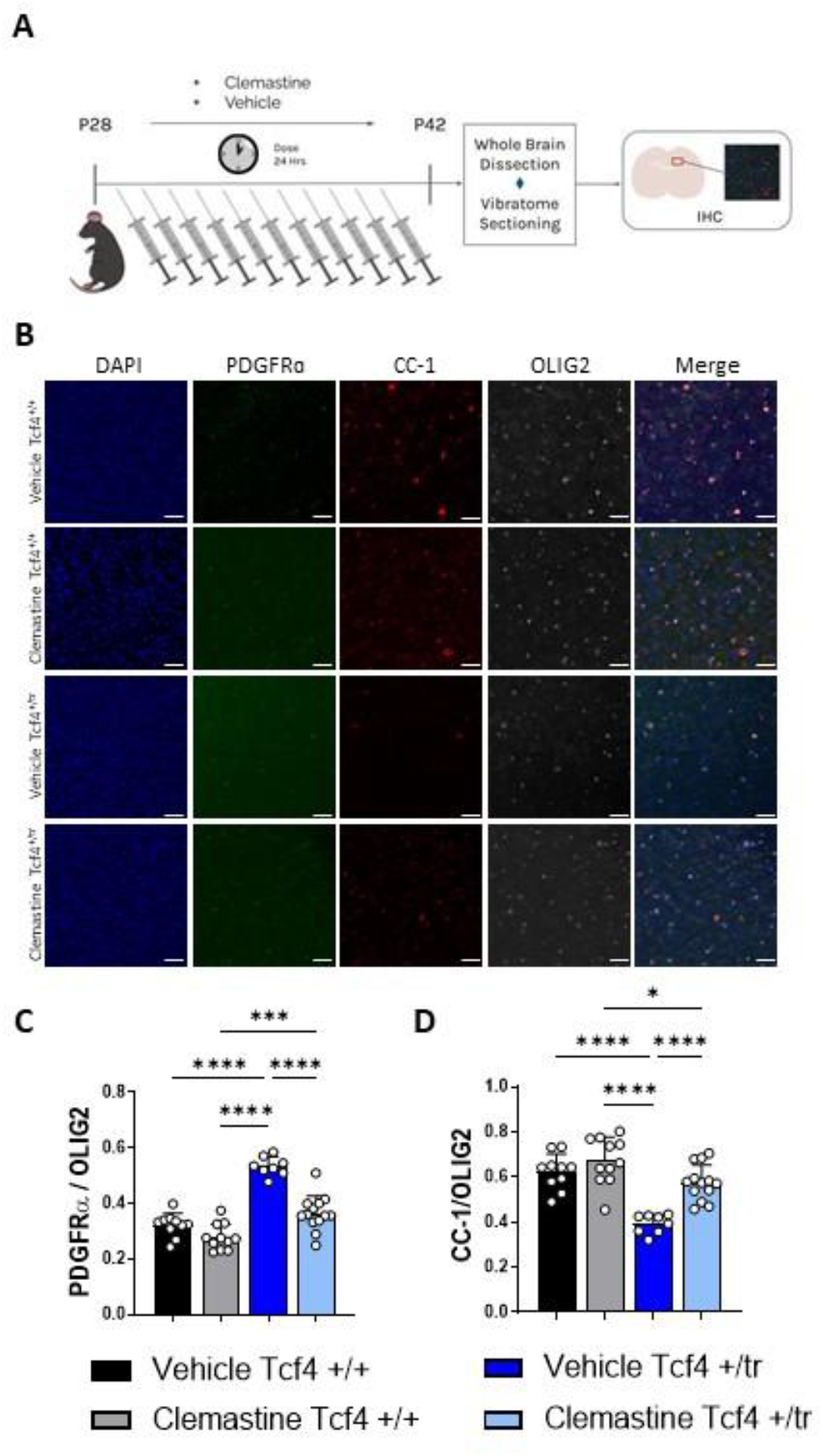
Clemastine increases maturation of oligodendrocytes in *Tcf4*^*+/tr*^ mice. **(A)** Graphical representation of the dosing regime in vivo. **(B)** Immunohistochemistry of OPC/OL in 55μM tissue slices following 14 days of either vehicle (DMSO) or clemastine administration. Tissue stained for DAPI (blue), PDGFRα (green), MBP (red) and Olig2 (gray). **(C-D)** Summary plots show processed tissue from *Tcf4*^*+/+*^ mice with vehicle (black) or treated with clemastine fumarate (gray) as well as tissue from *Tcf4*^*+/tr*^ mice with vehicle (dark blue) or treated with clemastine fumarate (sky blue). Representative results for response to vehicle and clemastine administration comparing PDGFRα/Olig2 (immature population) (Two-way ANOVA n=10-13 biologically independent animals/genotype F_3,23_=58.28 p<0.0001 data are presented as mean values ± SEM) and CC-1/Olig2+ (mature population) (Two-way ANOVA n=10-13 biologically independent animals/genotype F_3,23_=26.20 p<0.0001 data are presented as mean values ± SEM) These data show clemastine fumarate increased the percentage of mature OLs (MBP) while reducing the overall pool of OPCs (PDGFRα) in vivo. *p<0.05, **p<0.01, ^***p<0.001, ****^ p<0.0001.

### Clemastine increases myelination activity and overall myelination in PTHS mice

In order to visualize myelination, we used transmission electron microscopy (TEM) in both *Tcf4*^*+/+*^ and *Tcf4*^*+/tr*^ littermates that had been dosed with either vehicle or clemastine (Figure 4A). Blinded to genotype and condition, TEM images were taken on the CC directly above the hippocampus from anatomically equivalent tissue sections in littermates. Following blinded quantification, we replicated our previous observation that Tcf4 mutation leads to a significant reduction in myelinated axons in the corpus callosum (Figure 4B, C). However, we observed that the two-week clemastine treatment was unable to restore *Tcf4*^*+/+*^ levels of total myelin density in Tcf4^+/tr^ mice (Figure 4B, C). We next classified myelinated axons as either compacted or uncompacted and observed that the balance between myelinated (compacted) and ongoing myelination (uncompacted) was significantly different between vehicle-treated *Tcf4*^*+/tr*^ mice and vehicle-treated *Tcf4*^*+/+*^ littermates (Figure 4D, E), suggesting that the active process of myelination was reduced in Tcf4^+/tr^ mice. Notably, we observed that clemastine treatment significantly increased the density of uncompacted myelin in *Tcf4*^*+/tr*^ mice compared to vehicle-treated *Tcf4*^*+/t*r^ mice. Thus, normalizing the proportion of uncompacted myelin to levels observed in vehicle-treated *Tcf4*^*+/+*^ mice (Figure 4E), which indicates clemastine treatment is promoting the generation of new myelinated axons in the corpus callosum. Consistent with the in vivo and in vitro immunochemistry data, we note that clemastine is not only effective at enhancing the production of mature OLs (Figures 1-2), it also promotes the generation of new myelin ensheathment throughout this relatively short dosage time frame.

**Figure 3.**
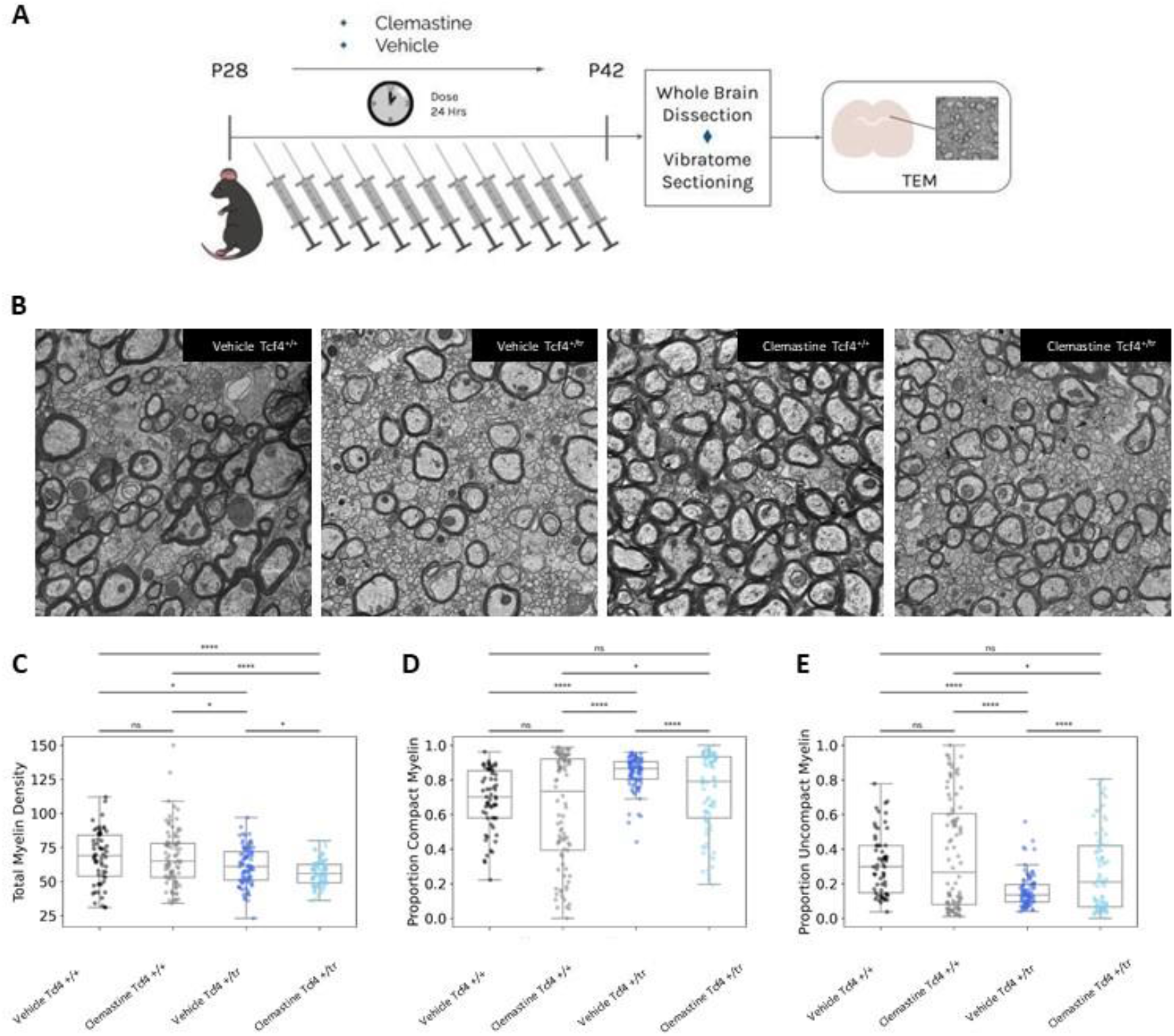
Clemastine increases the proportion of uncompacted myelin in *Tcf4*^*+/tr*^ mice. **(A)** Graphical representation of the dosing regime and subsequent TEM. **(B)** Representative electron micrographs of the CC from vehicle/clemastine *Tcf4*^*+/+*^ and vehicle/clemastine *Tcf4*^*+/tr*^ mice. Arrows indicate compact and uncompact myelin designations. **(C)** Density of myelination across all images for all conditions showing reduction of myelination in the vehicle and clemastine treated animals. (Two-way ANOVA biologically independent animals/genotype F=1.54, p=0.22) **(D)** Proportion of compact myelin per image showing less compaction overall in the clemastine Tcf4^+/+^ and clemastine *Tcf4*^*+/tr*^. (Two-way ANOVA biologically independent animals/genotype F=1.24, p-value=0.26) **(E)** Proportion of uncompact myelin per image showing clemastine treatment increases uncompact myelin in both *Tcf4*^*+/+*^ and *Tcf4*^*+/tr*^ mice. (Two-way ANOVA biologically independent animals/genotype F=1.24 p-value=0.26) Center values represent the mean and error bars are S.E.M., *p<0.05, **p<0.01, ***p<0.001, ****p<0.0001.

**Figure 4.**
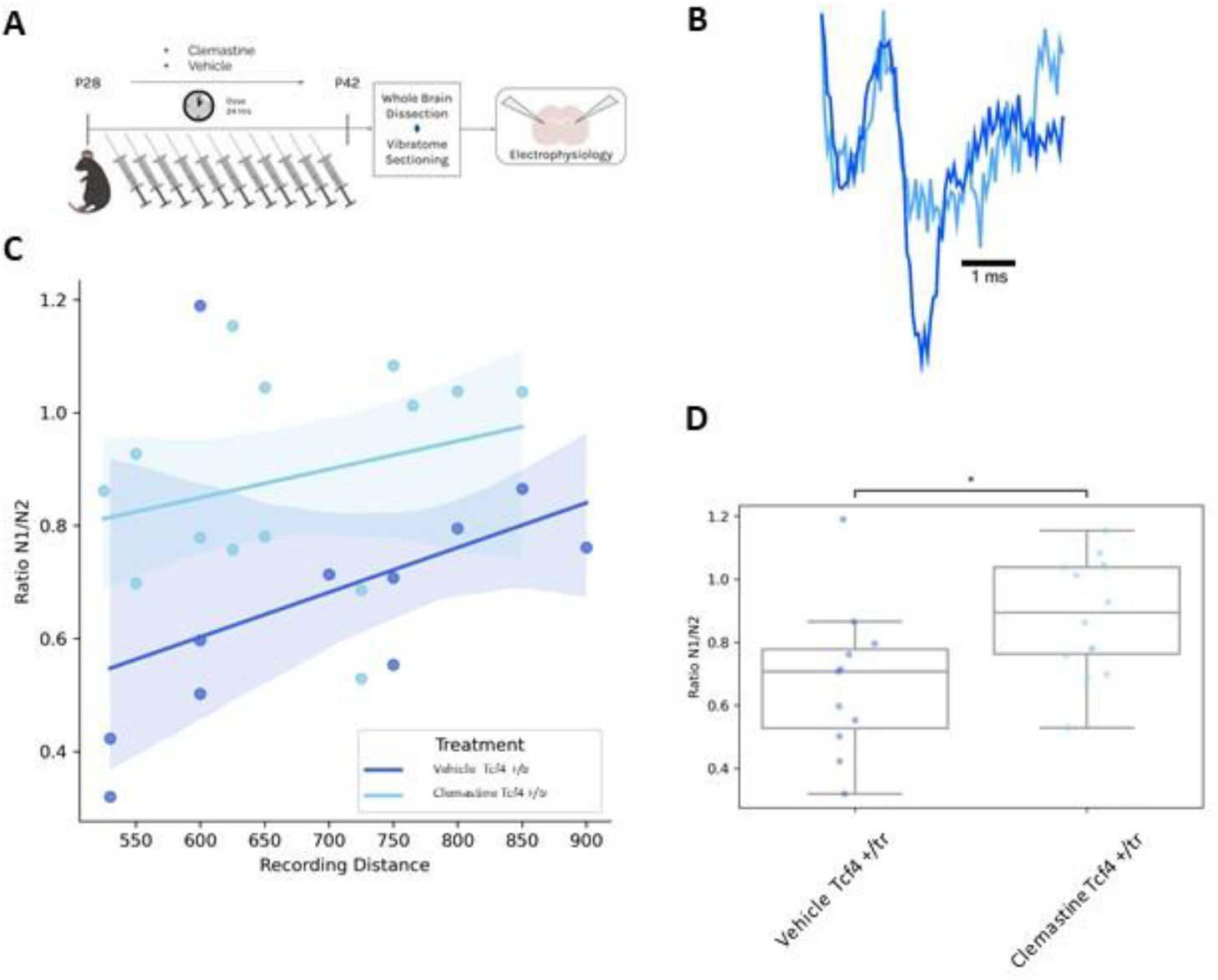
Clemastine rescues compound action potential phenotypes in acute brain slices from *Tcf4*^*+/tr*^ mice. **(A)** Graphical representation of the dosing regime and subsequent electrophysiology. **(B)** Representative electrophysiology traces of evoked compound action potentials recorded in the CC from vehicle *Tcf4*^*+/tr*^ and clemastine treated *Tcf4*^*+/tr*^ mice. N1 represents action potentials traveling down myelinated axons and N2 represents action potentials traveling down unmyelinated axons. Example traces are normalized by the N1 peak. **(C)** The proportion of action potentials traveling down myelinated axons was consistently reduced in vehicle *Tcf4*^*+/tr*^ mice compared to clemastine treated *Tcf4*^*+/tr*^ mice. (p=0.034) **(D)** Linear regression fit of the N1/N2 ratio at its respective recording distance (ANCOVA F=2.72, p-value=0.019).

### Clemastine rescues pathological reduction of myelinated axons in PTHS mice

We next determined if clemastine’s effect on the OL population was effective at normalizing physiology in *Tcf4*^*+/tr*^ mice. We measured the propagation of compound action potentials (CAPs) in the corpus callosum (CC) in acute brain slices from *Tcf4*^*+/+*^ and *Tcf4*^*+/tr*^ mice. CAPs were evoked by a bipolar stimulating electrode and recorded by a field electrode placed at varying distances across the CC and the amplitude of N1 and N2 peaks was quantified (Figure 4A). The N1 and N2 peaks represent CAPs traveling down myelinated and unmyelinated axons, respectively (22), and we previously showed the N1/N2 ratio was significantly reduced in *Tcf4*^*+/tr*^ mice (6). Two-week clemastine treatment of *Tcf4*^*+/tr*^ mice was effective at significantly increasing the N1/N2 ratio compared to vehicle-treated *Tcf4*^*+/tr*^ littermates (Figure 4C). These results indicate clemastine treatment is effective at improving the physiological function in the CC of the PTHS mouse model.

### Clemastine rescues behavioral deficits in PTHS mice

Given the clemastine-dependent increases in the mature OL population and rescued electrophysiological in *Tcf4*^+/tr^ mice, we were interested to determine if these changes were adequate to ameliorate behavioral deficits in these mice. It was previously shown that a variety of PTHS mouse models display consistent behavioral deficits, including hyperlocomotion and reduced anxiety in the open field among others (2,5,23). Therefore, we treated *Tcf4*^*+/tr*^ mice and *Tcf4*^*+/+*^ littermates with clemastine for 14 days and then assayed their behavior in the open field (Figure 5A). Untreated *Tcf4*^*+/tr*^ animals exhibited an overall greater distance traveled and increased frequency of entering the center of the field (inversely related to anxiety) (Figure B, C). Remarkably, clemastine-treated *Tcf4*^*+/tr*^ mice showed a significant reduction in total distance traveled and a reduction in the frequency of entering the center of the field, which was similar to untreated *Tcf4*^*+/+*^ mice (Figure 5B, C). These data indicate that clemastine treatment is effective at normalizing hyperlocomotion and anxiety phenotypes in the PTHS mouse model. All together, these data suggest remyelination therapy appears to be beneficial to cells, circuits, and behavior in a preclinical model of PTHS and supports the notion that clemastine or other remyelination agents could be an applicable therapeutic intervention in humans diagnosed with PTHS.

**Figure 5.**
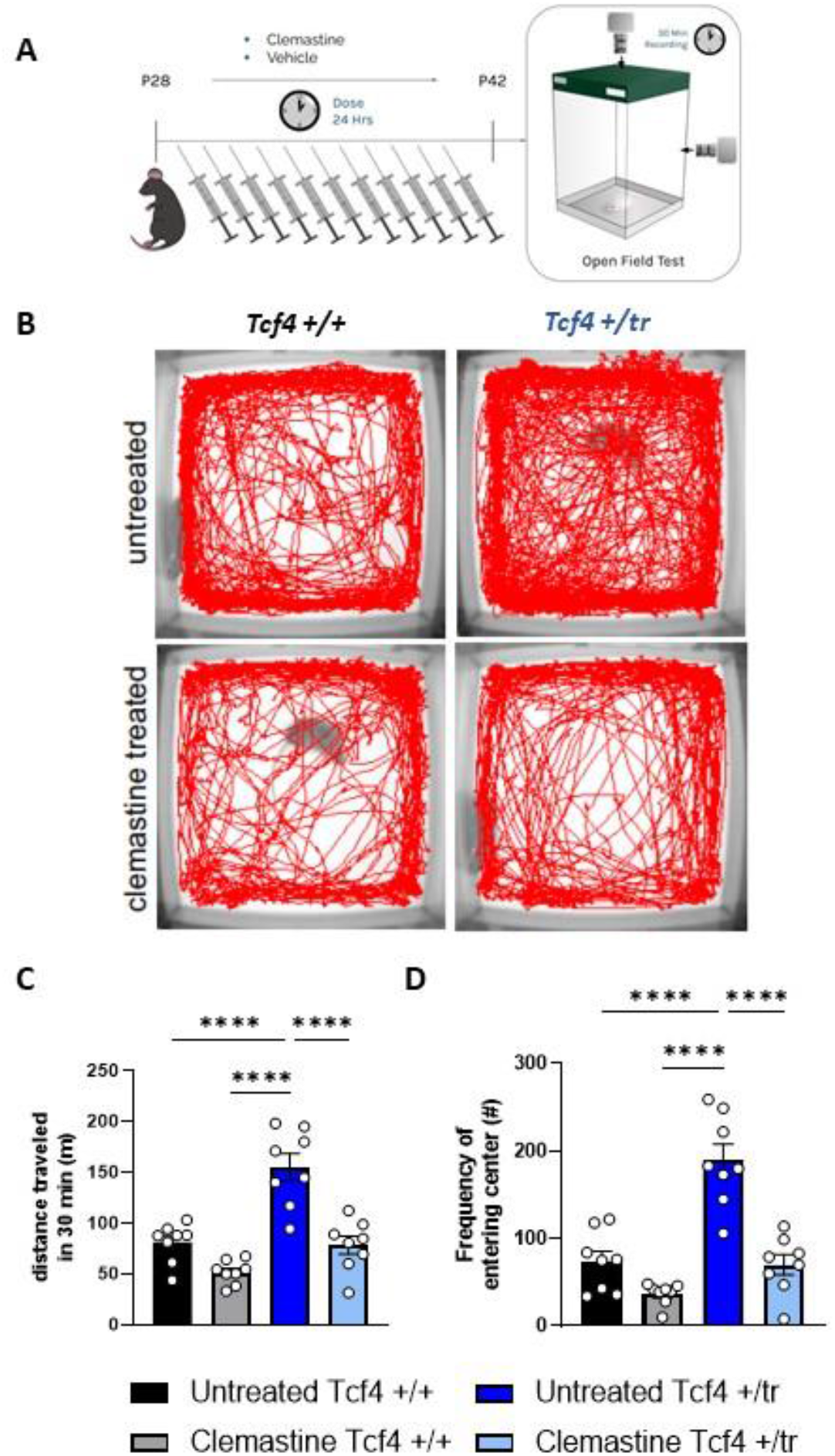
Clemastine rescues behavioral deficits in *Tcf4*^*+/tr*^ mice. **(A)** Locomotor activity maps from *Tcf4*^*+/+*^ and *Tcf4*^*+/tr*^ mice untreated or treated for 14 days with clemastine fumarate. Movement was recorded for 30 min following placement in a 37.5 cm x 37.5 cm novel open field arena. **(B)** Summary plots for distance traveled over 30 min depicts *Tcf4*^*+/+*^ mice untreated (black) or treated with clemastine fumarate (gray) as well as *Tcf4*^*+/tr*^ mice untreated (dark blue) or treated with clemastine fumarate (sky blue). These data show that *Tcf4*^*+/tr*^ mice treated with clemastine fumarate exhibited locomotor activity similar to treated *Tcf4*^*+/+*^ mice (*n* = 8 biologically independent animals/genotype, *T*_14_ = 0.2403, *p* > 0.05, data are presented as mean values ± SEM). **(C)** Summary plots for frequency of entering center (middle 15.25 cm x 15.25 cm) show that *Tcf4*^*+/tr*^ mice treated with clemastine fumarate exhibited a similar frequency of entering the center area compared to treated *Tcf4*^*+/+*^ mice (*n* = 8 biologically independent animals/genotype, *T*_14_ = 0.2081, *p* > 0.05, data are presented as mean values ± SEM). Asterisk (*) indicate the difference between mice groups (unpaired *t*-test). One symbol = *p* < 0.05, two symbols = *p* < 0.01, three symbols =*p* < 0.001, four symbols = *p* < 0.0001.

## Discussion

We have performed a series of experiments to demonstrate that pharmacological enhancement of myelination is effective at normalizing abnormal brain function and behavior in a mouse model of PTHS. We demonstrate that clemastine, a previously described remyelination compound, is effective at normalizing OPC and OL density both in vitro and in vivo (Figure 1,2). Additionally, EM analysis showed that clemastine treatment elevated the levels of uncompacted myelin indicative of an increase in ongoing myelination (Figure 3). This improved OL maturation resulted in functional rescue, whereby more CAPs were observed traveling down myelinated axons following treatment with clemastine (Figure 4). Lastly,we showed that the clemastine-dependent improvement of OL populations and physiological function normalized behavioral deficits (Figure 5). Together, these results provide strong evidence that remyelination may be a suitable therapeutic approach for PTHS.

### Myelination deficits in PTHS

Myelination deficits are consistently observed in several mouse models of PTHS. The initial observation was derived from transcriptomic analysis of five different PTHS mouse models and was subsequently biologically confirmed in the same *Tcf4*^*+/tr*^mouse model used in this study (6). Regulation of oligodendrocyte development by *Tcf4* was also shown in the mouse spinal cord where homozygous knockout of the long isoform of *Tcf4* resulted in a significant reduction in mature OLs (9). Moreover, conditional deletion of *Tcf4* in the *Nkx2*.*1* lineage resulted in a significant increase in OPCs of the mouse olfactory bulb (10). *Tcf4* is expressed in all stages of OL development as demonstrated by single cell sequencing and fluorescent in situ hybridization, and a cell autonomous effect of *Tcf4* on OL phenotypes is established (6,8–10). Clinical evidence for myelination deficits in PTHS patients is currently qualitative, due to the rare occurrence of this syndrome, with reports of several patients showing delayed myelination, white matter hyperintensities, and dysplasia of the corpus callosum (24–28). Future studies using PTHS patient-derived induced pluripotent stem cells will be important to demonstrate OL phenotypes observed in the mouse model translate to the human condition.

### Therapeutic Potential of Clemastine

The remyelinating capabilities of clemastine were first identified in a drug screen that was specific for functional myelination (12). Clemastine is an FDA-approved, first generation antihistamine, that readily crosses the blood brain barrier, however its effect on OPCs appears to be through its off-target antimuscarinic effects (12,16). Clemastine administration has rescued myelination defects in a variety of mouse models of demyelination, neurodegeneration, and a neurodevelopmental disorder (11,15–17,29–32). Moreover, it showed remyelination capabilities in a phase 2 clinical trial for multiple sclerosis (33). Here we demonstrate that clemastine is effective at rescuing myelination deficits and behavior in a mouse model of PTHS. Consistent with previous findings, our results suggest the cellular mechanism for clemastine’s effect is through promoting OPCs to differentiate, as we showed both in vitro and in vivo that clemastine shifts the OL population by reducing OPCs and increasing OL density (Figure 1,2). Clemastine’s effect on OPC differentiation is proposed to work through antagonism of the M1R (12–14). To our knowledge,there are no reports of dysregulation of M1R function downstream of *Tcf4* mutations and *Chrm1* expression is not altered in PTHS mouse models (6). Therefore, it is likely that the molecular mechanism of rescue by clemastine is through an indirect mechanism that is not related to dysregulated pathways downstream of the *Tcf4* mutation. If this is the case, our data would suggest that a variety of remyelination therapeutics have the potential to normalize myelination in the PTHS mouse model because there is no requirement that the remyelination treatment directly normalize *Tcf4*-dependent transcription.

### Myelination deficits in ASD

Evidence for myelination defects across the autism spectrum is building and remyelination therapies could be a potential therapeutic target when deficits in myelination are suspected. We previously found that convergent differentially expressed genes (DEGs) across three syndromic ASD mouse models (*Tcf4, Mecp2*, and *Pten*) were enriched for biological terms related to oligodendrocytes and myelination (6). Importantly, the eigengene of these overlapping DEGs could be used to separate ASD cases from controls in a large sample of postmortem ASD brains. Moreover, cellular deconvolution of these same postmortem ASD samples predicted an overall reduction in OL fraction of RNA, suggesting there may be a reduction in OLs in idiopathic ASD (6). Re-analysis of single cell sequencing data from 41 postmortem ASD and control brains also identified a reduction in OL abundance (6,34). In addition, many identified ASD risk genes are known to regulate development and function of cells in the OL lineage. For instance, *SCN2a*, which is considered a high confidence syndromic ASD gene (35,36) is expressed in OPCs and regulates their excitability and development (37,38). Other high confidence ASD risk genes such as CHD8 (39–42), ADNP (43), PTEN (44–46), and POGZ (47) to name a few, have also been shown to regulate the OL lineage and/or display abnormal myelination in the patient population. The process of myelination temporally coincides with progression of ASD and immediately precedes the first appearances of the disorder (18). In addition, it was shown that ongoing myelination is a requirement for long-term consolidation of spatial memory and motor learning in several mouse models (48–50), which suggests abnormal developmental myelination could underlie intellectual disability and motor deficits, which are common comorbidities in ASD. Fortunately, OPCs are the most abundant proliferating cell population in the human brain and are present throughout the lifespan, which makes them suitable therapeutic targets for NDDs. All together these studies strongly support the concept of remyelination as a potential treatment for not only PTHS, but potentially many subtypes of NDDs and ASDs which are predicted or shown to have myelination delays or deficits.

## Figure Legends

**Table 1.** List of Antibodies.

| **Antibody** | **Dilution** | **Company** | **Catalog No.** |
| --- | --- | --- | --- |
| Rabbit anti-Olig2 | 1:500 | EMD Millipore | AB9610 |
| Mouse anti-APC (CC-1) | 1:500 | Calbiochem | OP80-100UG |
| Rat anti-PDGFRα | 1:500 | BD Pharmingen | 558774 |
| Mouse anti-MBP (F-6) | 1:500 | Santa Cruz Biotechnology | Sc-271524 |
| Rat anti-MBP (aa82-87) | 1:50 | Bio Rad | MCA409S |
| Alexa Fluor 546 Goat anti-Rabbit | 1:1000 | Invitrogen | A11010 |
| Alexa Fluor 488 Goat anti-Rat | 1:1000 | Invitrogen | A48262 |
| Alexa Fluor 647 Donkey anti-Mouse | 1:1000 | Invitrogen | A31571 |

## Acknowledgements

We are grateful for the vision and generosity of the Lieber and Maltz families, who made this work possible. We thank the Johns Hopkins School of Medicine Microscope Core Facility and specifically LaToya Roker and Michael Delannoy for generating TEM images of CC used in this study. This work was supported by the Lieber Institute, the Pitt-Hopkins Research Foundation Award (B.J.M.), a NIMH grant R01MH110487 (B.J.M.). The content is solely the responsibility of the authors and does not necessarily represent the official views of the National Institutes of Health.

## Author Contributions

J.F.B. performed ICC and confocal imaging; J.F.B., S.R.S., G.S., N.L.S. and R.F.K. performed IHC; J.F.B., and D.D. performed EM experiments, B.J.M. performed electrophysiology experiments; C.M.C. and D.K.M. performed behavior experiments; I.P.B. formulated compounds and contributed to dosing design; J.F.B., S.R.S, N.L.S. and B.J.M. contributed to experimental design, data analyses and writing. All authors discussed the results and edited the manuscript.

## References

1. Chen H-Y, Bohlen JF, Maher BJ. Molecular and Cellular Function of Transcription Factor 4 in Pitt-Hopkins Syndrome. Dev Neurosci. 2021 Jun 16;43(3–4):159–67.

2. Thaxton C, Kloth AD, Clark EP, Moy SS, Chitwood RA, Philpot BD. Common Pathophysiology in Multiple Mouse Models of Pitt-Hopkins Syndrome. J Neurosci. 2018 Jan 24;38(4):918–36.

3. Cleary CM, James S, Maher BJ, Mulkey DK. Disordered breathing in a Pitt-Hopkins syndrome model involves Phox2b-expressing parafacial neurons and aberrant Nav1.8 expression. Nat Commun. 2021 Oct 13;12(1):5962.

4. Grubišić V, Kennedy AJ, Sweatt JD, Parpura V. Pitt-Hopkins Mouse Model has Altered Particular Gastrointestinal Transits In Vivo. Autism Res. 2015 Oct;8(5):629–33.

5. Kennedy AJ, Rahn EJ, Paulukaitis BS, Savell KE, Kordasiewicz HB, Wang J, et al. Tcf4 regulates synaptic plasticity,DNA methylation, and memory function. Cell Rep. 2016 Sep 6;16(10):2666–85.

6. Phan BN, Bohlen JF, Davis BA, Ye Z, Chen H-Y, Mayfield B, et al. A myelin-related transcriptomic profile is shared by Pitt-Hopkins syndrome models and human autism spectrum disorder. Nat Neurosci. 2020 Mar;23(3):375–85.

7. Kim H, Berens NC, Ochandarena NE, Philpot BD. Region and cell type distribution of TCF4 in the postnatal mouse brain. Front Neuroanat. 2020 Jul 17;14:42.

8. Marques S, Zeisel A, Codeluppi S, van Bruggen D, Mendanha Falcão A, Xiao L, et al. Oligodendrocyte heterogeneity in the mouse juvenile and adult central nervous system. Science. 2016 Jun 10;352(6291):1326–9.

9. Wedel M, Fröb F, Elsesser O, Wittmann M-T, Lie DC, Reis A, et al. Transcription factor Tcf4 is the preferred heterodimerization partner for Olig2 in oligodendrocytes and required for differentiation. Nucleic Acids Res. 2020 May 21;48(9):4839–57.

10. Zhang X, Huang N, Xiao L, Wang F, Li T. Replenishing the aged brains: targeting oligodendrocytes and myelination? Front Aging Neurosci. 2021 Nov 25;13:760200.

11. Lee JI, Park JW, Lee KJ, Lee DH. Clemastine improves electrophysiologic and histomorphometric changes through promoting myelin repair in a murine model of compression neuropathy. Sci Rep. 2021 Oct 22;11(1):20886.

12. Mei F, Fancy SPJ, Shen Y-AA, Niu J, Zhao C, Presley B, et al. Micropillar arrays as a high-throughput screening platform for therapeutics in multiple sclerosis. Nat Med.2014 Aug;20(8):954–60.

13. Chen J-F, Liu K, Hu B, Li R-R, Xin W, Chen H, et al. Enhancing myelin renewal reverses cognitive dysfunction in a murine model of Alzheimer’s disease. Neuron. 2021 Jul 21;109(14):2292–2307.e5.

14. Minigh J. Clemastine. xPharm: The Comprehensive Pharmacology Reference. Elsevier; 2008. p. 1–6.

15. Li Z, He Y, Fan S, Sun B. Clemastine rescues behavioral changes and enhances remyelination in the cuprizone mouse model of demyelination. Neurosci Bull.2015 Oct;31(5):617–25.

16. Mei F, Lehmann-Horn K, Shen Y-AA, Rankin KA, Stebbins KJ, Lorrain DS, et al. Accelerated remyelination during inflammatory demyelination prevents axonal loss and improves functional recovery. eLife. 2016 Sep 27;5.

17. Liu J, Dupree JL, Gacias M, Frawley R, Sikder T, Naik P, et al. Clemastine enhances myelination in the prefrontal cortex and rescues behavioral changes in socially isolated mice. J Neurosci. 2016 Jan 20;36(3):957–62.

18. Deoni SCL, Zinkstok JR, Daly E, Ecker C, MRC AIMS Consortium, Williams SCR, et al. White-matter relaxation time and myelin water fraction differences in young adults with autism. Psychol Med. 2015 Mar;45(4):795–805.

19. Yoshida A, Takashima K, Shimonaga T, Kadokura M, Nagase S, Koda S. Establishment of a simple one-step method for oligodendrocyte progenitor cell preparation from rodent brains. J Neurosci Methods. 2020 Aug 1;342:108798.

20. Maher BJ, LoTurco JJ. Disrupted-in-schizophrenia (DISC1) functions presynaptically at glutamatergic synapses. PLoS ONE. 2012 Mar 30;7(3):e34053.

21. Mickelsen LE, Bolisetty M,Chimileski BR, Fujita A, Beltrami EJ, Costanzo JT, et al. Single-cell transcriptomic analysis of the lateral hypothalamic area reveals molecularly distinct populations of inhibitory and excitatory neurons. Nat Neurosci. 2019 Apr;22(4):642–56.

22. Olmos-Serrano JL, Kang HJ, Tyler WA, Silbereis JC, Cheng F, Zhu Y, et al. Down syndrome developmental brain transcriptome reveals defective oligodendrocyte differentiation and myelination. Neuron. 2016 Mar 16;89(6):1208–22.

23. Ekins S,Puhl AC,Davidow A. Repurposing the dihydropyridine calcium channel inhibitor nicardipine as a nav1.8 inhibitor in vivo for pitt hopkins syndrome. Pharm Res. 2020 Jun 11;37(7):127.

24. Amiel J, Rio M, de Pontual L,Redon R, Malan V, Boddaert N, et al. Mutations in TCF4, encoding a class I basic helix-loop-helix transcription factor, are responsible for Pitt-Hopkins syndrome, a severe epileptic encephalopathy associated with autonomic dysfunction. Am J Hum Genet. 2007 May;80(5):988–93.

25. Goodspeed K, Newsom C, Morris MA, Powell C, Evans P, Golla S. Pitt-Hopkins Syndrome: A Review of Current Literature, Clinical Approach, and 23-Patient Case Series. J Child Neurol. 2018 Mar;33(3):233–44.

26. Rosenfeld JA, Leppig K, Ballif BC, Thiese H, Erdie-Lalena C, Bawle E, et al. Genotype-phenotype analysis of TCF4 mutations causing Pitt-Hopkins syndrome shows increased seizure activity with missense mutations. Genet Med. 2009 Nov;11(11):797–805.

27. Brockschmidt A, Filippi A, Charbel Issa P, Nelles M, Urbach H, Eter N, et al. Neurologic and ocular phenotype in Pitt-Hopkins syndrome and a zebrafish model. Hum Genet. 2011 Nov;130(5):645–55.

28. Stavropoulos DJ, MacGregor DL, Yoon G. Mosaic microdeletion 18q21 as a cause of mental retardation. Eur J Med Genet. 2010 Dec;53(6):396–9.

29. Xie Y-Y, Pan T-T, Xu D-E, Huang X, Tang Y, Huang W, et al. Clemastine ameliorates myelin deficits via preventing senescence of oligodendrocytes precursor cells in alzheimer’s disease model mouse. Front Cell Dev Biol. 2021 Oct 21;9:733945.

30. Barak B, Zhang Z, Liu Y, Nir A, Trangle SS, Ennis M, et al. Neuronal deletion of Gtf2i, associated with Williams syndrome, causes behavioral and myelin alterations rescuable by a remyelinating drug. Nat Neurosci. 2019 May;22(5):700–8.

31. Deshmukh VA, Tardif V, Lyssiotis CA, Green CC, Kerman B, Kim HJ, et al. A regenerative approach to the treatment of multiple sclerosis. Nature. 2013 Oct 17;502(7471):327–32.

32. Cree BAC, Niu J, Hoi KK, Zhao C, Caganap SD, Henry RG, et al. Clemastine rescues myelination defects and promotes functional recovery in hypoxic brain injury. Brain. 2018 Jan 1;141(1):85–98.

33. Green AJ, Gelfand JM, Cree BA, Bevan C, Boscardin WJ, Mei F, et al. Clemastine fumarate as a remyelinating therapy for multiple sclerosis (ReBUILD): a randomised, controlled, double-blind, crossover trial. Lancet. 2017 Dec 2;390(10111):2481–9.

34. Velmeshev D,Schirmer L, Jung D, Haeussler M, Perez Y,Mayer S,et al. Single-cell genomics identifies cell type-specific molecular changes in autism. Science.2019 May 17;364(6441):685–9.

35. Fu JM, Satterstrom FK,Peng M, Brand H,Collins RL,Dong S, et al. Rare coding variation illuminates the allelic architecture, risk genes, cellular expression patterns, and phenotypic context of autism. medRxiv. 2021 Dec 21;

36. Satterstrom FK, Kosmicki JA, Wang J, Breen MS, DeRubeis S, An J-Y, et al. Large-Scale Exome Sequencing Study Implicates Both Developmental and Functional Changes in the Neurobiology of Autism. Cell. 2020 Feb 6;180(3):568–584.e23.

37. Berret E, Barron T, Xu J, Debner E, Kim EJ, Kim JH. Oligodendroglial excitability mediated by glutamatergic inputs and Nav1.2 activation. Nat Commun. 2017 Sep 15;8(1):557.

38. Gould E, Kim JH. SCN2A contributes to oligodendroglia excitability and development in the mammalian brain. Cell Rep. 2021 Sep 7;36(10):109653.

39. Marie C, Clavairoly A, Frah M, Hmidan H, Yan J, Zhao C, et al. Oligodendrocyte precursor survival and differentiation requires chromatin remodeling by Chd7 and Chd8. Proc Natl Acad Sci USA. 2018 Aug 28;115(35):E8246–55.

40. Jung H, Park H, Choi Y, Kang H, Lee E, Kweon H,et al. Sexually dimorphic behavior, neuronal activity, and gene expression in Chd8-mutant mice. Nat Neurosci. 2018 Sep;21(9):1218–28.

41. Zhao C, Dong C, Frah M, Deng Y, Marie C, Zhang F, et al. Dual requirement of CHD8 for chromatin landscape establishment and histone methyltransferase recruitment to promote CNS myelination and repair. Dev Cell. 2018 Jun 18;45(6):753–768.e8.

42. Kawamura A, Katayama Y, Nishiyama M, Shoji H, Tokuoka K, Ueta Y, et al. Oligodendrocyte dysfunction due to Chd8 mutation gives rise to behavioral deficits in mice. Hum Mol Genet. 2020 May 28;29(8):1274–91.

43. Van Dijck A, Vulto-van Silfhout AT, Cappuyns E, van der Werf IM, Mancini GM, Tzschach A, et al. Clinical presentation of a complex neurodevelopmental disorder caused by mutations in ADNP. Biol Psychiatry. 2019 Feb;85(4):287–97.

44. Tilot AK, Gaugler MK, Yu Q, Romigh T, Yu W, Miller RH, et al. Germline disruption of Pten localization causes enhanced sex-dependent social motivation and increased glial production. Hum Mol Genet. 2014 Jun 15;23(12):3212–27.

45. Frazier TW, Embacher R, Tilot AK, Koenig K, Mester J, Eng C. Molecular and phenotypic aabnormalities in individuals with germline heterozygous PTEN mutations and autism. Mol Psychiatry. 2015 Sep;20(9):1132–8.

46. Goebbels S, Oltrogge JH, Kemper R, Heilmann I, Bormuth I, Wolfer S, et al. Elevated phosphatidylinositol 3,4,5-trisphosphate in glia triggers cell-autonomous membrane wrapping and myelination. J Neurosci. 2010 Jun 30;30(26):8953–64.

47. Suliman-Lavie R, Title B, Cohen Y, Hamada N, Tal M, Tal N, et al. Pogz deficiency leads to transcription dysregulation and impaired cerebellar activity underlying autism-like behavior in mice. Nat Commun.2020 Nov 17;11(1):5836.

48. Steadman PE, Xia F, Ahmed M, Mocle AJ,Penning ARA, Geraghty AC, et al. Disruption of oligodendrogenesis impairs memory consolidation in adult mice. Neuron. 2020 Jan 8;105(1):150–164.e6.

49. McKenzie IA, Ohayon D, Li H, de Faria JP, Emery B, Tohyama K, et al. Motor skill learning requires active central myelination. Science. 2014 Oct 17;346(6207):318–22.

50. Pan Y, Cai K, Cheng M, Zou X, Li M. Responsive Social Smile: A Machine Learning based Multimodal Behavior Assessment Framework towards Early Stage Autism Screening. 2020 25th International Conference on Pattern Recognition (ICPR). IEEE; 2021. p. 2240–7.

